# Characterization of high affinity IgM and IgG monoclonal antibodies against norovirus variants GII.4 and GII.17

**DOI:** 10.1101/2024.05.11.593658

**Authors:** Jumpei Tagawa, Saeko Yanaka, Yuri Kato, Akitsu Masuda, Jae Man Lee, Akinobu Senoo, Kosuke Oyama, Motohiro Nishida, Takahiro Kusakabe, Jose M.M. Caaveiro

**Affiliations:** Graduate School of Pharmaceutical Sciences, Kyushu University, 3-1-1 Maidashi, Higashi, Fukuoka 812-8582, Japan; Labolatory for Materials and Structures, Institute of Innovative research, Tokyo Institute of Technology, 4259 Nagatsuda, Midori-ku, Yokoyama, Kanagawa 226-8503, Japan; 3Laboratory of Creative Science for Insect Industries, Kyushu University Graduate School of Bioresource and Bioenvironmental Sciences, 744 Motooka, Nishi-ku, Fukuoka, 819-0395, Japan; Laboratory of Insect Genome Science, Kyushu University Graduate School of Bioresource and Bioenvironmental Sciences, 744 Motooka, Nishi-ku, Fukuoka, 819-0395, Japan

**Author notes:** Corresponding author: Jose M.M. Caaveiro.

## Abstract

Human norovirus, a leading cause of viral gastroenteritis, results in significant global health and economic burden, requiring sensitive and accurate diagnosis and effective therapeutics and vaccines. In this study, we immunized mice with the virus like capsid particles of GII.4, a mainstream strain, and GII.17, a modern strain that began to circulate in 2014, and used hybridoma technology to generate hybridoma cells that produce norovirus-binding antibodies against GII.4 and GII.17, respectively. Selection of these hybridoma cells yielded monoclonal IgG and IgM antibodies against these strains. Characterization of these antibodies revealed that avidity effect by multivalent binding is necessary for IgM to bind to norovirus at high efficiency, while IgG achieve high affinity even by monovalent binding. Surface plasmon resonance and ELISA data suggest that the high density of antigen protrusion domain in the norovirus capsid, containing approximately protomers, facilitates IgM to bind to norovirus capsid with high efficiency.

## Introduction

Human norovirus is recognized as the leading cause of sporadic and epidemic viral gastroenteritis (1, 2) and is estimated to be responsible for approximately 699 million norovirus infections, more than 1 million hospitalizations, and 219,000 deaths worldwide, resulting in a social cost of $60 billion annually (3, 4). Thus, the global burden caused by norovirus is substantial and requires sensitive and accurate diagnosis and effective therapeutics and vaccination.

Based on the primary sequence of VP1 protein of the capsid, noroviruses have been phylogenetically classified into at least 10 gene groups (GI-GX) and further subdivided into 49 genotypes (e.g., GII.4) (5). Among the norovirus genotypes, GII.4 strains represent the dominant group of human noroviruses worldwide (55-85%) and has been the predominant genotype for more than 20 years (6, 7). Significant attention has thus been focused on the genotype of GII.4 strains in human populations (8, 9). However, the emergence of a new genotype, strain GII.17, in the winter of 2014-2015, caught the world’s attention and surpassed strain GII.4 as the main cause of norovirus outbreaks in several countries in late 2014 (10, 11). Genomic nucleotide sequences of structural proteins can differ by more than 50% between gene groups (12), and antibodies produced by some GII.4 strains lack cross-reactivity to GII.17 strains (13, 14). Because genetic diversity in structural proteins also causes changes in antigenic properties, it is essential to understand the human immune response to infection via human noroviruses and antigenic variation among circulating human norovirus strains.

Noroviruses are non-enveloped RNA viruses whose outer surface is covered by a capsid protein composed of 180 monomeric units. The components of the capsid are the capsid protein VP1 and the minor structural protein VP2 (15); VP1 comproses two domains, a highly conserved shell (S) domain and a more variable protruding (P) domain (16). In the human immune system, infection with noroviruses results in the production of neutralizing antibodies against these components.

The complexity of human norovirus cross-reactivity and neutralization by human antibodies is not yet fully realized. A number of studies have evaluated the existence of a human polyclonal immune response to human norovirus (17–19). Studies analyzing mAbs isolated from norovirus-infected humans and mice using hybridoma technology have primarily isolated IgG-type antibodies and have also identified IgA and IgM-type antibodies (20–23). Previous using epitope analysis of norovirus antibodies have identified neutralizing epitopes on norovirus capsids primarily against the protruding 2 (P2) domain, but also conserved epitopes in inaccessible regions of the viral capsid (24, 25). Indeed, antigen mapping studies using strain-specific P and S domains suggested that some of the highly cross-reactive mAbs bind to the S domain (26, 27). Epitope analysis of IgGs against GII.4 strains has advanced because of the variety of their antibodies. In contrast, epitope mapping of mAbs against GII.17 is limited because of the few examples of GII.17-specific mAbs (28, 29). Moreover, detailed studies of anti-norovirus antibodies of the IgM class are even more limited. Because VLP are complex entities with high density of P-domains, which is the major epitope of norovirus antibodies, IgM displaying 10 or 12 antigen-binding domains could bind with high affinity due to their multivalency (30–32).

In this study, we obtained hybridoma cells producing monoclonal antibodies that specifically recognized strains GII.4 or GII.17. Antibodies of the IgG and IgM class were obtained, and the molecular basis of their efficient binding to norovirus evaluated by complementary techniques.

## Results and Discussion

### Selection of hybridoma cells secreting antibodies against norovirus

Hybridoma cells were generated in triplicate, thus dividing the selection of hybridoma cells into three groups. Hybridomas producing antibodies specific to GII.4 and GII.17 were selected by a two-step selection process. The first and second groups of hybridoma cells were selected by ELISA using norovirus VLPs, and the third group was selected by ELISA using norovirus P-domain (Figure 1A).

**Fig.1.**
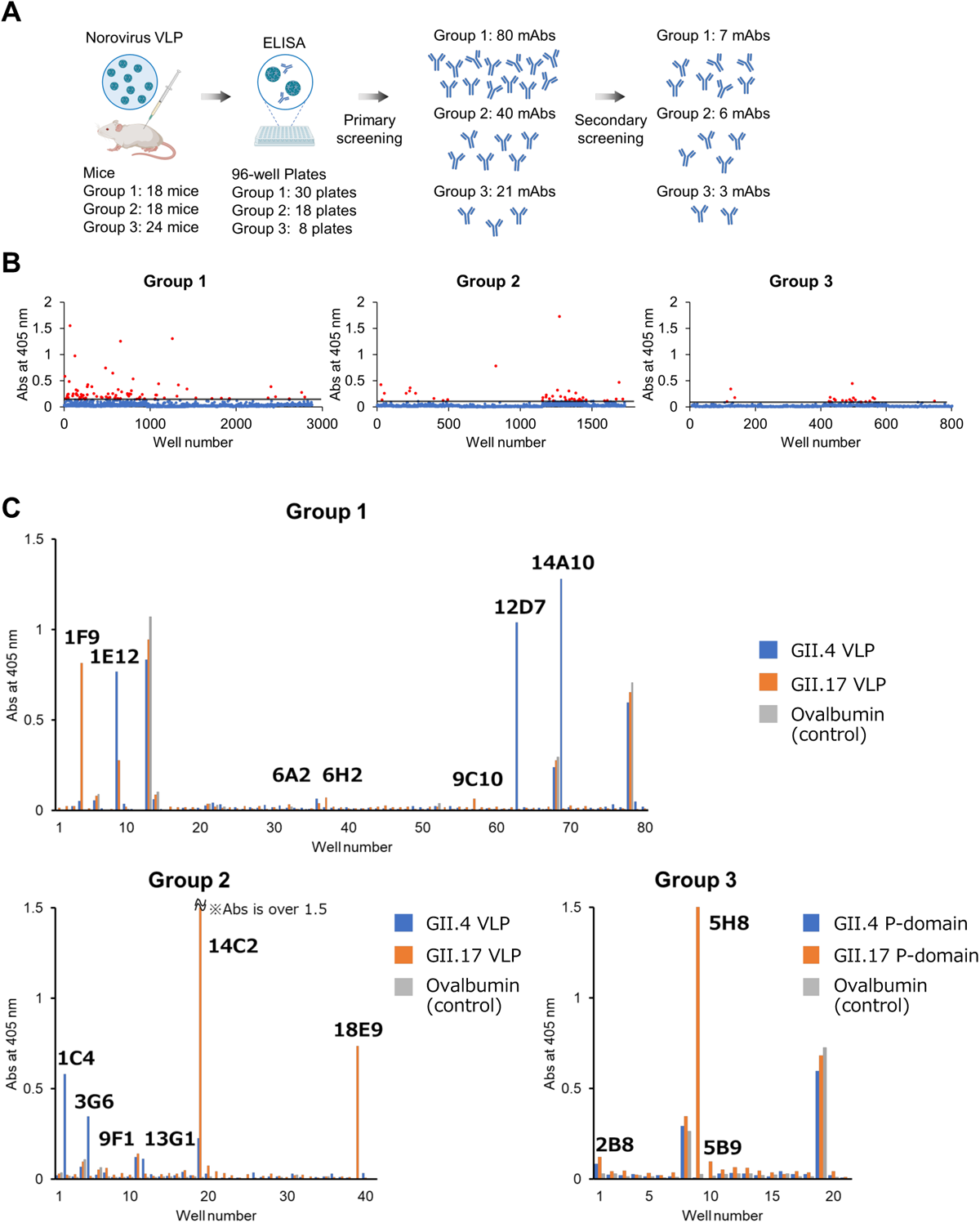
Selection of hybridoma cells producing anti-norovirus antibodies. (A) Mice immunized with norovirus VLPs were sacrificed and plasma cells from the spleens of immunized mice were fused with myeloma cells to create hybridoma cells expressing mAbs. Cells producing Abs that reacted with wells decorated with norovirus protein were identified by ELISA. A total of 141 wells identified in the primary screening. Secondary screening also employing ELISA identified 16 mAbs, which excluded Abs with unspecific binding and false positives from the primary screening. Samples belonging to groups 1 and 2 were evaluated against norovirus VLPs, whereas samples belonging to group 3 were selected against the P-domain. (B) ELISA corresponding to the primary screening; red indicates clones that were used for further analysis. (C) ELISA for secondary screening, with the names of clones used for further analysis.

In the primary screening, hybridoma cells generated from spleen cells derived from 60 mice (group 1, group 2 and group 3 consisted of 18, 18 and 24 animals, respectively), were seeded to 56 96-well plates (group 1, group 2 and group 3 corresponded to 30, 18 and 8 plates, respectively). Hybridoma cells from 141 different wells (group 1, group 2, and group 3 gave 80, 40, and 21 positives, respectively). ELISA positives corresponded to antigens containing the sequence of the GII.4 strain or to the sequence of the GII.17 strain (Figure 1B).

In the secondary screening, additional ELISA was performed with the hybridoma cells obtained from the 141 wells to exclude nonspecific binding. To that end we employed the norovirus unrelated protein ovalbumin. After this step 16 different samples of hybridoma cells were selected. These cells corresponded to group 1, group 2 and group 3 containing 7, 6, and 3 positives, respectively. The 16 hybridoma cell samples were monoclonalized by the method of ultra-dilution yielding seven types of hybridoma cells that we termed according to their position in the plate (12D7, 14A10, 3G6, 13G1, 14C2, 18E9, and 5H8) (Figure 1C).

### Identification of antibody isotypes and their purification

Antibody classes and subclasses were determined for the seven different mAbs-producing hybridoma cells using a commercial kit (Table 1). Of the seven antibodies examined, five were found to be of the IgM class and two of the IgG class. To purify the IgG-class antibodies (14C2 and 5H8), we employed Protein G affinity chromatography. For the IgM antibodies (which are not retained by protein A or protein G) we had to consider alternative methods for purification. Protein L is known to specifically bind to the light chain with a known sequence. We conducted sequence analysis of the light chains of the obtained IgM and revealed that sequence motif important for protein L binding are preserved for two of them (12D7 and 3G6). For theses IgM, protein L was used for purification. The other IgMs were purified using an IgM affinity column, but with lower specificity towards IgM than ProteinA or ProteinL have for IgG. Therefore, 13G1 and 18E9 showed slightly lower purity in the SDS-PAGE (Figure 2B). Secondary purification was then performed using size exclusion chromatography (SEC), confirming elution peaks at around 150 kDa for antibodies of the IgG class, and peaks around 900 kDa for the antibodies of the IgM class (Figure 2A). Analysis by SDS-PAGE under reducing conditions confirmed the purification of the IgG-type antibody and IgM-type antibody.

**Table. 1.**
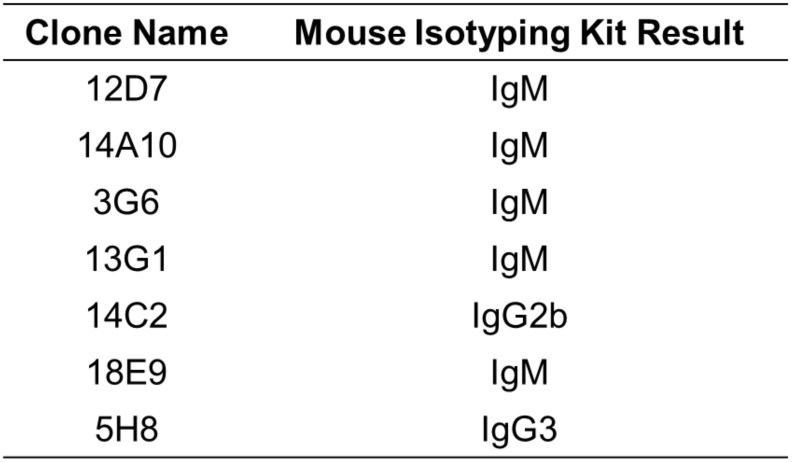
Antibody isotypes.

| Clone Name | Mouse Isotyping Kit Result |
| --- | --- |
| 12D7 | IgM |
| 14A10 | IgM |
| 3G6 | IgM |
| 13G1 | IgM |
| 14C2 | IgG2b |
| 18E9 | IgM |
| 5H8 | IgG3 |

**Fig. 2.**
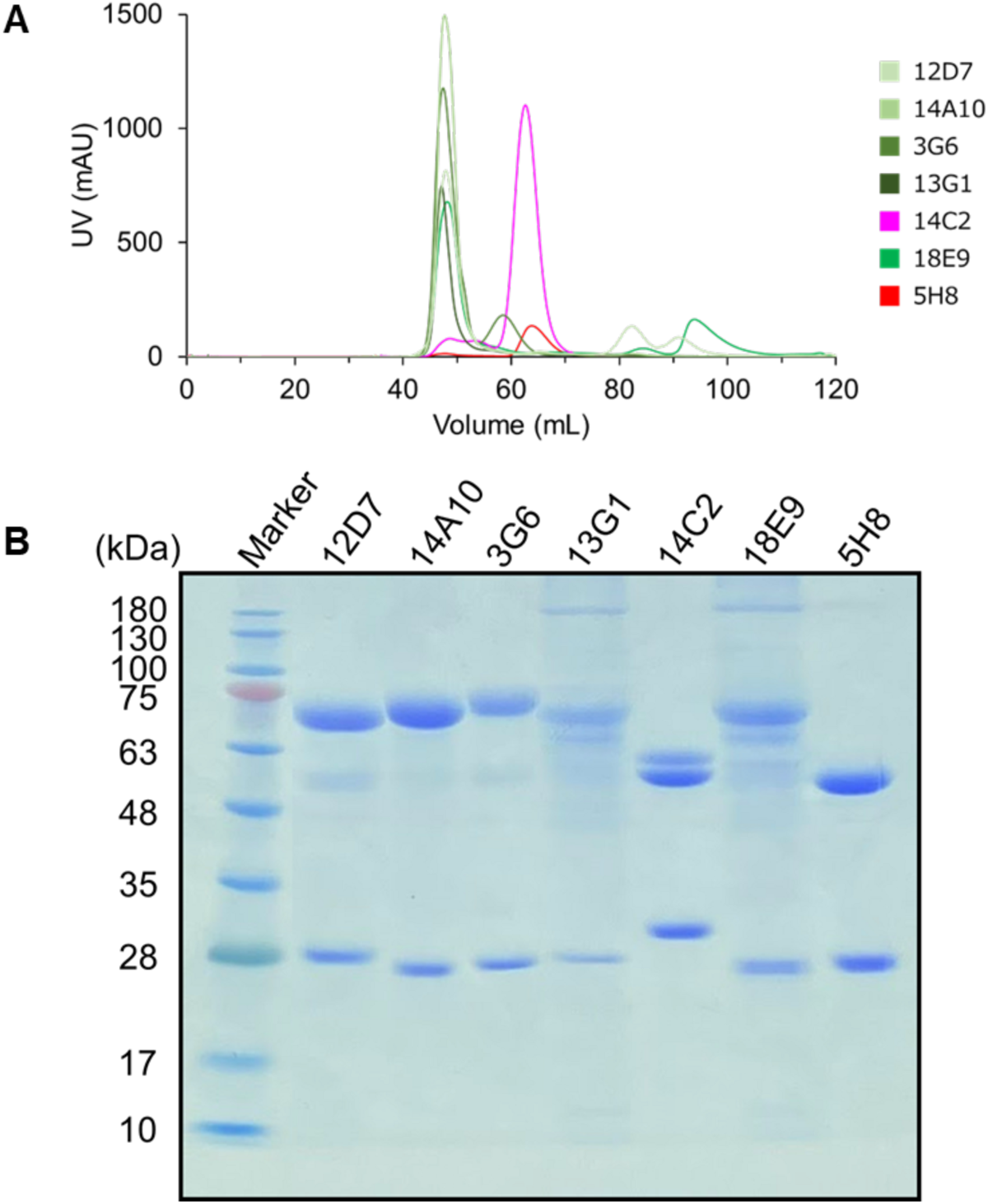
Purification of anti-norovirus monoclonal antibodies. (A) Overlaid chromatograms from size exclusion chromatography of mAbs, with IgM-type antibodies shown in green and IgG-type antibodies in red. (B) Reduced SDS-PAGE analysis of purified mAbs.

### Determination of binding affinity of mAbs to norovirus P-domain by surface plasmon resonance (SPR)

The binding affinities of the seven monoclonal antibodies obtained to the P-domains of GII.4 and GII.17 strains of norovirus were determined by surface plasmon resonance (SPR) (Table 2). The isolated antibodies could be classified into three groups based on their binding specificity to norovirus P-domains. The first group (GII.4) is a group that binds only to the norovirus projection domain of GII.4 strains and includes two mAbs, 12D7 and 14A10, both of which are IgM-type antibodies (Figure 3). High affinity in on the nM order was observed for 12D7 and 14A10 when P-domains of GII.4 was immobilized on the SPR surface. In contrast, no binding was observed for IgM antibodies 12D7 and 14A10 when the IgMs were immobilized and P-domain of GII.4 was used as analyte (Table 3). Therefore, it is suggested that IgM antibodies 12D7 and 14A10 can bind to antigens with the avidity effect with multiple antigen-binding sites, although each Fab have weak antigen-binding ability. The second group (GII.17), which binds only to the norovirus P-domain of GII.17, includes two antobodies, 14C2 and 5H8, both of which are IgG-type antibodies. The binding affinity of 14C2 and 5H8 was very high in the order of pM when the P-domain of GII.17 was immobilized on the SPR surface. When the antibodies were immobilized and the P-domain of GII.17 were flowed, 14C2 showed a 1000-fold decrease in affinity, whereas 5H8 showed almost no decrease in affinity. These data indicated that 14C2, an IgG-type antibody, bind to P-domain of GII.17 bivalently with clear signs of avidity effect, whereas 5H8 binds to the P-domain of GII.17 monovalently. The third group antibodies (3G6, 13G1, and 18E9) did not show significant binding to either the norovirus P-domain of GII.4 or GII.17 when evaluated by SPR, although they showed significant binding in the ELISA during screening.

**Table. 2.** Kinetic analysis for the interaction of anti-norovirus antibodies with immobilized P-domain.

| Variant specificity | Subclass | Clone name (Ligand) | Variant name (Analyte) | $K_{on}$ (/Ms) | $K_{off}$ (/s) | $K_D$ (M) |
| --- | --- | --- | --- | --- | --- | --- |
| GII.4 | IgM | 12D7 | GII.4 | $1.85 \times 10^6$ | $1.71 \times 10^{-2}$ | $9.21 \times 10^{-9}$ |
|  |  |  | GII.17 | N.D | N.D | N.D |
| | IgM | 14A10 | GII.4 | $3.31 \times 10^4$ | $1.06 \times 10^{-5}$ | $3.21 \times 10^{-10}$ |
|  |  |  | GII.17 | N.D | N.D | N.D |
| GII.17 | IgG | 14C2 | GII.4 | N.D | N.D | N.D |
| | | | GII.17 | $1.16 \times 10^5$ | $1.07 \times 10^{-7}$ | $9.21 \times 10^{-13}$ |
|  | IgG | 5H8 | GII.4 | N.D | N.D | N.D |
| | | | GII.17 | $2.11 \times 10^5$ | $1.09 \times 10^{-5}$ | $5.15 \times 10^{-11}$ |

**Table. 3.** Kinetic analysis for the interaction of P-domain with immobilized anti-norovirus antibodies.

| Variant specificity | Subclass | Variant name (Ligand) | Clone name (Analyte) | $K_{on}$ (/Ms) | $K_{off}$ (/s) | $K_D$ (M) |
| --- | --- | --- | --- | --- | --- | --- |
| GII.4 | IgM | GII.4 | 12D7 | N.D | N.D | N.D |
|  |  | GII.17 |  | - | - | - |
|  | IgM | GII.4 | 14A10 | N.D | N.D | N.D |
|  |  | GII.17 |  | - | - | - |
| GII.17 | IgG | GII.4 | 14C2 | - | - | - |
| | | GII.17 | | $8.29 \times 10^4$ | $1.29 \times 10^{-4}$ | $1.55 \times 10^{-9}$ |
|  | IgG | GII.4 | 5H8 | - | - | - |
| | | GII.17 | | $7.83 \times 10^5$ | $6.37 \times 10^{-5}$ | $8.13 \times 10^{-11}$ |

**Fig. 3.**
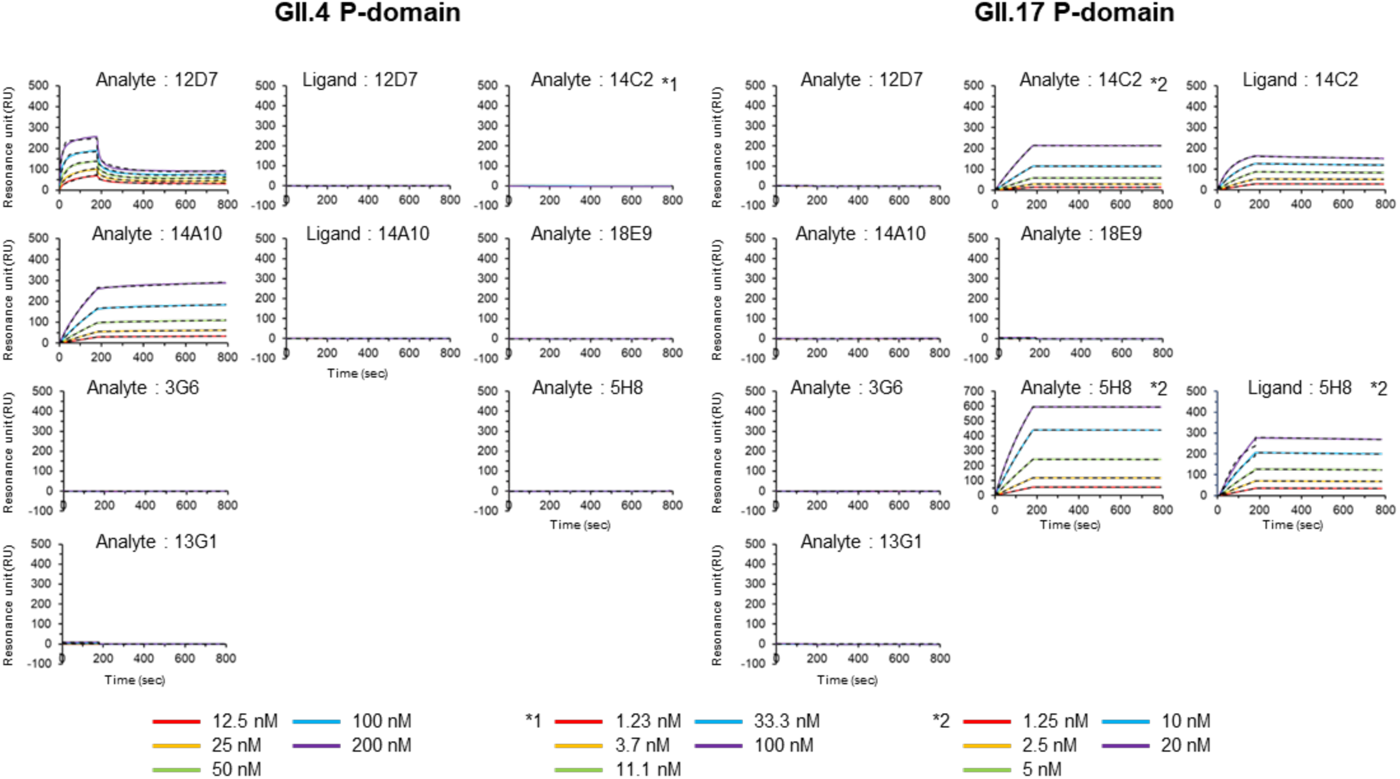
Binding of anti-norovirus monoclonal antibodies to the capsid P-domain. Binding of mAbs to norovirus P-domain was evaluated by SPR. Sensorgrams of binding to GII.4 strains are shown on the left and those to GII.17 strains on the right. The solid line shows the obtained sensorgrams and the fitting is shown as a dashed line.

We speculate that this discrepancy is caused by the difference in the density (number) of antigen molecules in each assay. Norovirus VLPs have a capsid structure comprising 180 units of the P-domain, and therefore a large number of molecules of P-domain at short distance from each other are present in the VLP particles employed in the ELISA selection step. In contrast, when norovirus P-domain monomers are immobilized on ELISA plates uniformly, the distance between each antigen is expected to grow with respect to that in the VLP particles immobilized on the ELISA plate. In the second screening ELISA, we immobilized GII.4 VLP, which is a particle highly packed with very high density of P-domains of GII.4. In the SPR experiment, the density of molecules of P-domain of GII.4 immobilized as monomers is lower than that of P-domain molecules in the VLP. Indeed, the data show that 12D7 and 14A10 suggests that avidity effect is extremely important for the binding of IgM antibodies. Therefore, we suspected that the density of antigen difference affected the binding of IgM antibodies. As SPR were not applicable for detection of the binding of very large VLP particles, we examined this possibility using ELISA.

### Examination of multivalent binding of IgMs by ELISA

To further analyze the binding specificity of the seven mAbs to VLPs vs the P-domain of noroviruses, binding assays were performed using ELISA. As we have expected, 3G6, 13G1, and 18E9 showed binding only to GII.4 VLP and not to P-domains of GII.4 (Figure 4), supporting our idea that density of the antigen affects the binding of IgM antibodies. Furthermore, the binding profile of antibodies that bind to both the VLP and the projection domain of norovirus were found to be significantly different between IgG-type and IgM-type antibodies. 12D7 and 14A10, IgM-type antibodies, showed an absorbance of about one eighth that of P-domain in relation to that of the VLP. The absorbance of the P-domain of IgG antibodies 14C2 and 5H8 showed little decrease compared to the absorbance of the VLP. This also supports the hypothesis that high affinity of IgM is largely dependent on the avidity effect of multivalent binding.

**Fig. 4.**
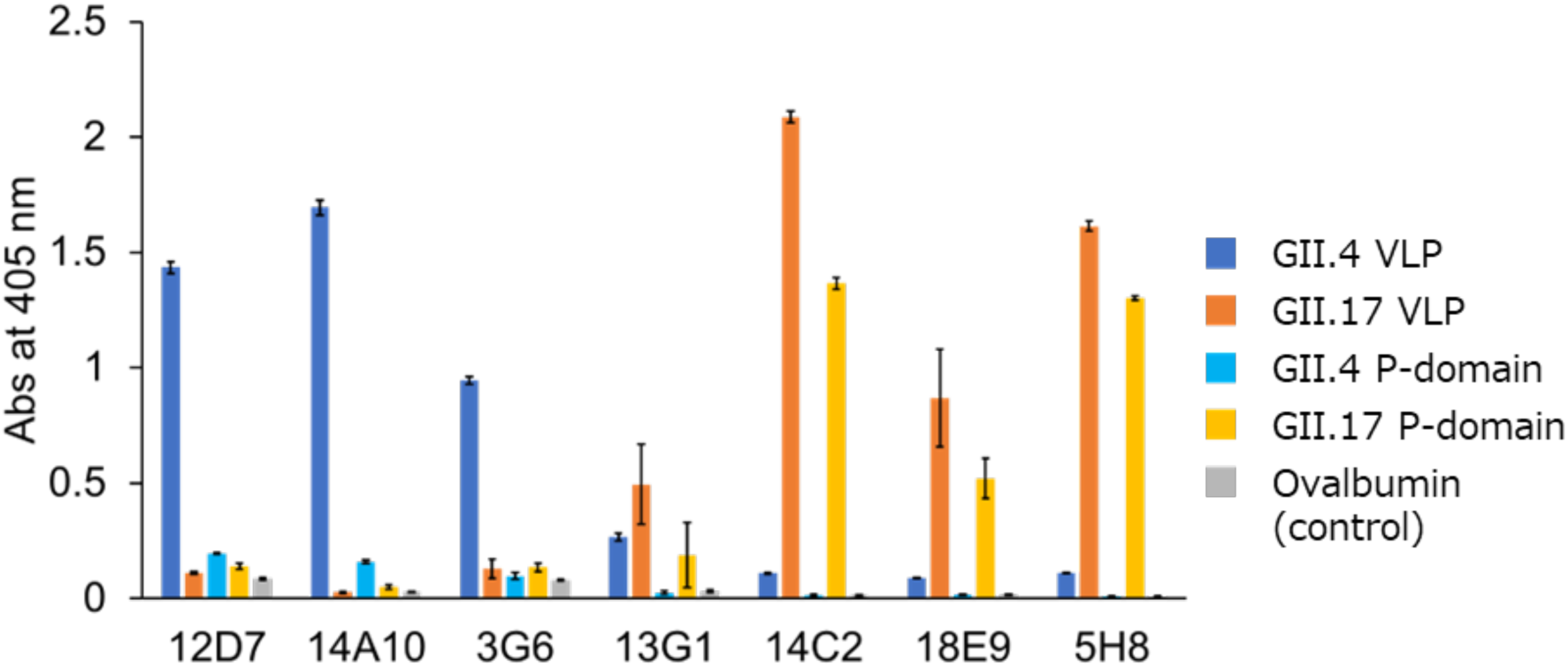
Evaluation of binding specificity of anti-norovirus antibodies by ELISA. Indirect ELISA was used to evaluate the binding of seven mouse mAbs to GII.4 VLP, GII.17 VLP, GII.4 P-domain, and GII.17 P-domain, respectively. Error bars indicate standard deviation. (n=3)

The low immobilization density of the P-domain per unit area on the ELISA plate relative to the norovirus VLPs may have prevented the IgM-type antibodies from binding at high affinity, resulting in much lower absorbance in the ELISA relative to the VLPs. On the other hand, IgG-type antibodies bind bivalently to the antigen, but their monovalent binding is stronger than that of IgM-type antibodies and does not depend on the avidity effect to a large extent, which may explain why the absorbance of the ELISA was not much lower than that of the VLPs. In particular, for 5H8, the absorbance in ELISA was almost the same for VLPs and the P-domain, similar to the trend observed in SPR, suggesting that monovalent binding is very strong. Similar to the SPR results, only four antibodies (12D7, 14A10, 14C2, and 5H8) bind to the P-domain in ELISA, suggesting that 3G6, 13G1, and 18E9 bind to the shell domain rather than the P-domain.

We also speculated that the density difference of VLPs and norovirus P-domains affected the screening step. In group 1 and 2, VLP was used for selection and norovirus P-domain was used in group 3. IgMs were obtained in group 1 and 2 but not in group 3. This is reasonable since IgM has a greater tendency to bind to VLP with avidity effect rather than to the monomeric P-domain.

## Conclusion

In this study, we immunized mice with norovirus VLPs of the GII.4 strain (mainstream) infecting humans, and GII.17 strain (more recent strain that began to circulate in 2014), and following hybridoma technology and selection obtained anti-norovirus monoclonal antibodies against capsid proteins from GII.4 and GII.17 strains. Interaction analysis of the anti-norovirus antibodies obtained herein showed that they bind specifically to the capsid protein with high affinity. Most of the antibodies obtained were found to be of the IgM type, indicating that IgM efficiently interacts with antigens in the high-density, epitope-rich environment on the surface of VLPs.

## Materials and methods

### Preparation of mouse monoclonal antibodies to norovirus VLPs

Splenocytes from mice immunized with norovirus VLP were isolated and fused with HAT-sensitive mouse myeloma cells SP2/0 by the polyethylene glycol (PEG) method. Hybridoma cells producing monoclonal antibodies against norovirus VLPs were cloned by the limiting dilution method. Established hybridoma cells were cultured in 10% FBS E-RDF medium.

Hybridoma cells fused with spleen cells obtained from immunized mice and SP2/0 myeloma cells were seeded in 56 plates and hybridoma monoclones were obtained after 2 weeks. The mAbs produced by these clones were tested multiple times in an enzyme-linked immunosorbent assay (ELISA) against norovirus.

### ELISA screening and analysis

Immunoplates (Clear Flat-Bottom Immuno Nonsterile 96-Well Pates, Thermo Fisher Scientific) were incubated in coating buffer (15 mM Na_2_CO_3_, 35 mM NaHCO_3_, 3 mM NaN_3_, pH 9.6) (100 μL/well) coated with 1 μg/well concentrations of norovirus VLP GII.4 strain, norovirus VLP GII.17 strain, norovirus P-domain GII.4 strain, norovirus P-domain GII.17 strain and ovalbumin. Coated plates were sealed and incubated at 4°C overnight, washed three times with 0.05% PBS/Tween-20 (PBST) solution, and blocked with 300 μL/well of blocking buffer (PBS containing 5% skim milk) for 1 hour at 37°C. The plates were then washed three times with PBST and in ELISA screening, incubated with 100 μL of 2-fold dilution of hybridoma cell culture supernatant in PBS, while in ELISA analysis, incubated with 100 μL of serially diluted concentrations of each anti-norovirus antibody at 37°C for 1 hour. The optimal concentrations of anti-norovirus antibodies were normalized before testing binding ability and tested at concentrations beginning at 10 μg/well and then diluted serially. Next, the plates were washed three times with PBST, and horseradish peroxidase-conjugated goat anti-mouse IgG (H+L) was added as a secondary antibody and incubated at 37°C for 1 hour. Finally, the plate was washed three times with PBST, 60 μL of ABTS chromogenic agent was added, and 30 minutes later the OD value was measured at 405 nm.

### Expression and purification of norovirus P-domains

To determine the specificity of anti-norovirus binding antibodies, P1 and P2 domain sequences of norovirus GII.4 and GII.17 strains were recombinantly expressed. The sequences of the P-domain were cloned into a His-tagged pMAL-c6T vector. The P-domains were then expressed in *Escherichia coli* BL-21 cells and purified using Ni-NTA Agarose (FUJIFILM) and column chromatography.

### Expression and purification of IgG and IgM mAbs

Hybridoma cell lines producing Abs were gradually expanded from 96-well plates to 24-well plates, 6-well plates, 10 mm dishes, and finally to three 150 mm dishes per cell line. E-RDF medium at 37°C, 5% CO2 for 2 weeks, and the supernatant was used to determine antibody subclasses by Mouse Isotyping Kit (ICLLAB). The monoclonal antibodies produced were subjected to primary purification using HiTrap Protein G HP Columns, HiTrap Protein L Columns, and HiTrap IgM Purification HP Columns. IgG-type antibodies were purified using HiTrap Protein G HP Columns (Cytiva). The columns were operated at a constant flow rate of 1 ml/min. The column was equilibrated with 10 mL of binding buffer (20 mM sodium phosphate, pH 7.0), the supernatant containing IgG was applied to the column and the column was washed with 10 ml of binding buffer. Elution of bound IgG was performed in a one-step gradient using 0.1 M glycine buffer, pH 2.7, and 0.5 mL of the eluted fraction was collected. The elution fractions were immediately adjusted to physiological pH by adding 30 μl of 1 M Tris-HCl, pH 9.0. IgM-type antibodies 12D7 and 3G6 were purified using HiTrap Protein L Columns (Cytiva) and 14A10, 13G1, and 18E9 using HiTrap IgM Purification HP Columns.

For purification using HiTrap Protein L Columns, the column was operated at a constant flow rate of 1 ml/min. The column was equilibrated with 10 ml of binding buffer (20 mM sodium phosphate, 150 mM NaCl, pH 7.2); the supernatant containing IgM was applied to the column and the column was washed with 10 ml of binding buffer. Elution of bound IgM was performed in a one-step gradient with 0.1 M sodium citrate, pH 2.7) and 0.5 mL of the eluted fraction was collected. The elution fraction was immediately adjusted to physiological pH by adding 70 μl of 1 M Tris-HCl, pH 9.0. For purification using the HiTrap IgM Purification HP Column, ammonium sulfate was first added to the supernatant containing IgM to a final concentration of 0.8 M. To avoid precipitation of IgM, the sample was supplemented with a small amount of individual ammonium sulfate while continuously expanding the sample. The sample was passed through a 0.80 μm filter. The column was operated at a constant flow rate of 1 ml/min. The column was equilibrated with 10 ml of binding buffer (20 mM sodium phosphate, 0.8 M ammonium sulfate, pH 7.5). supernatant containing IgM was applied to the column and the column was washed with 10 ml of binding buffer. Elution of bound IgG was performed in a one-step gradient with 20 mM sodium phosphate, pH 7.5, and 0.5 mL of the eluted fraction was collected. Antibodies 12D7, 14A10, 13G1 14C2, 18E9, and 5H8 were purified using HiLoad 16/600 Superdex 200 pg and 3G6 was purified using HiLoad 16/600 Superdex 75 pg after primary purification by column chromatography.

### Binding analysis by SPR

SPR analysis was performed using a Biacore 8K system (GE Healthcare). Purified norovirus P-domain from GII.4 and GII.17 strains were immobilized to a CM5 sensor chip using the amine coupling method. 0.005% PBS/Tween20 (PBST) was used as reaction buffer, 0.005% PBS/Tween 20 (PBST) was used as the reaction buffer, and the flow rate was 30 μL/min. The affinity constant (*K*_*D*_) is equal to the ratio of the rate constants (*K*_*D*_ = *k*_*d*_ / *k*_*a*_, association rate constant (*k*_*a*_), dissociation rate constant (*k*_*d*_) and the antigen-antibody binds in a 1:1 kinetic model. To measure binding to norovirus P-domain GII.4 and GII.17 strains, the anti-norovirus binding antibody produced was serially diluted 3-fold to 1.23 nM or 2-fold to 25 nM in 0.005% PBS/Tween 20 (PBST) at a rate of 30 μL/min for 180 seconds. Dissociation was allowed to proceed for 600 seconds. The chip surface was regenerated with MgCl_2_ at a concentration of 3 M.

---

**Supplementary Figure 1.**
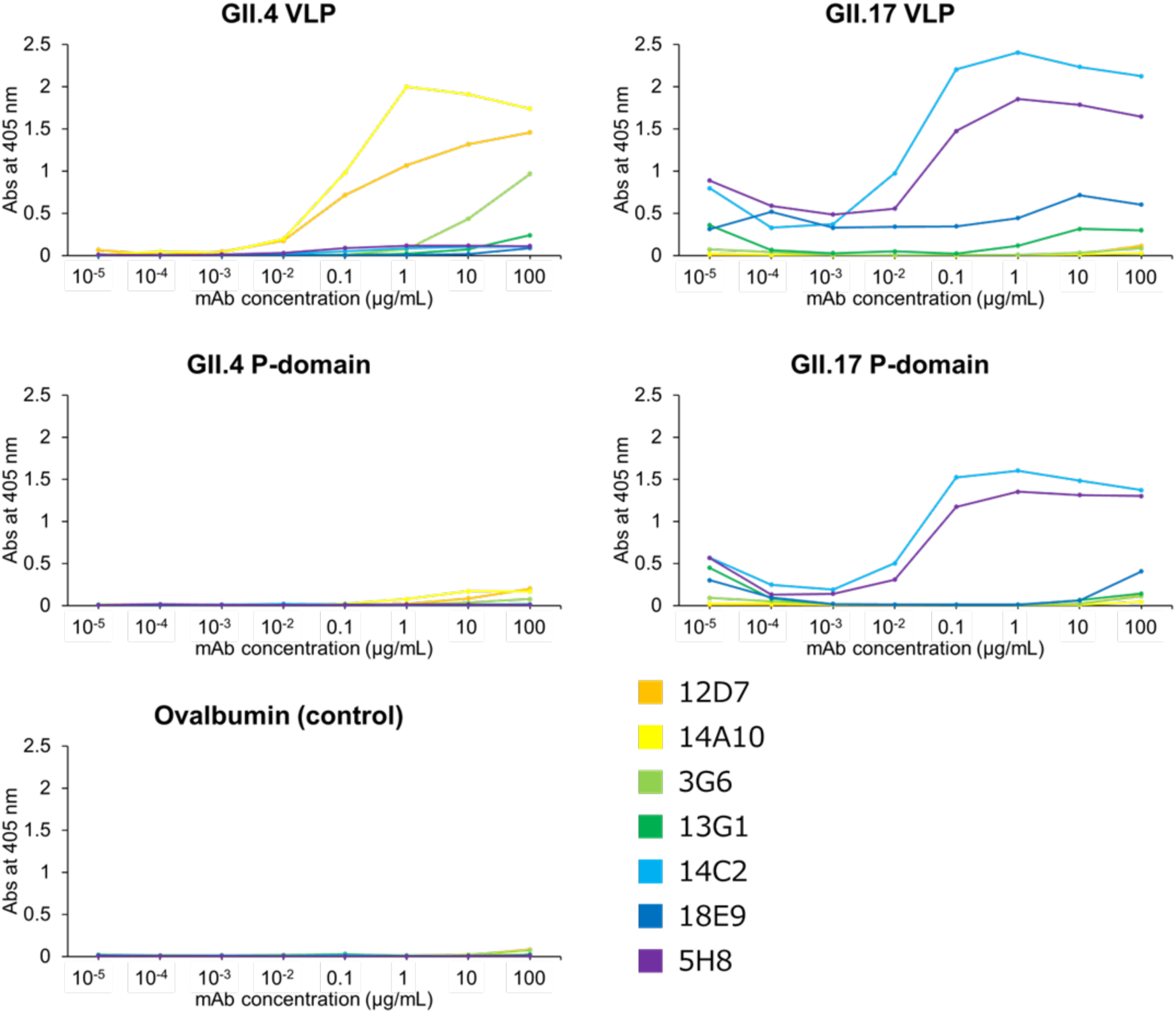
Evaluation of binding specificity of anti-norovirus antibodies by serial dilution ELISA. Indirect ELISA was used to evaluate binding of seven mouse mAbs to GII.4 VLP, GII.17 VLP, GII.4 P-domain, and GII.17 P-domain.

## References

1. Robilotti, E., Deresinski, S., and Pinsky, B. A. (2015) Norovirus. Clin Microbiol Rev. 28, 134–164

2. Patel, M. M., Widdowson, M.-A., Glass, R. I., Akazawa, K., Vinjé, J., and Parashar, U. D. (2008) Systematic Literature Review of Role of Noroviruses in Sporadic Gastroenteritis. Emerg. Infect. Dis. 14, 1224–1231

3. Bányai, K., Estes, M. K., Martella, V., and Parashar, U. D. (2018) Viral gastroenteritis. The Lancet. 392, 175–186

4. Bartsch, S. M., Lopman, B. A., Ozawa, S., Hall, A. J., and Lee, B. Y. (2016) Global Economic Burden of Norovirus Gastroenteritis. PLoS ONE. 11, e0151219

5. Chhabra, P., De Graaf, M., Parra, G. I., Chan, M. C.-W., Green, K., Martella, V., Wang, Q., White, P. A., Katayama, K., Vennema, H., Koopmans, M. P. G., and Vinjé, J. (2019) Updated classification of norovirus genogroups and genotypes. Journal of General Virology. 100, 1393–1406

6. Desai, R., Hembree, C. D., Handel, A., Matthews, J. E., Dickey, B. W., McDonald, S., Hall, A. J., Parashar, U. D., Leon, J. S., and Lopman, B. (2012) Severe Outcomes Are Associated With Genogroup 2 Genotype 4 Norovirus Outbreaks: A Systematic Literature Review. Clinical Infectious Diseases. 55, 189–193

7. Cannon, J. L., Bonifacio, J., Bucardo, F., Buesa, J., Bruggink, L., Chan, M. C.-W., Fumian, T. M., Giri, S., Gonzalez, M. D., Hewitt, J., Lin, J.-H., Mans, J., Muñoz, C., Pan, C.-Y., Pang, X.-L., Pietsch, C., Rahman, M., Sakon, N., Selvarangan, R., Browne, H., Barclay, L., and Vinjé, J. (2021) Global Trends in Norovirus Genotype Distribution among Children with Acute Gastroenteritis. Emerg. Infect. Dis. 27, 1438–1445

8. Lindesmith, L. C., Donaldson, E. F., LoBue, A. D., Cannon, J. L., Zheng, D.-P., Vinje, J., and Baric, R. S. (2008) Mechanisms of GII.4 Norovirus Persistence in Human Populations. PLoS Med. 5, e31

9. Tan, M., and Jiang, X. (2005) Norovirus and its histo-blood group antigen receptors: an answer to a historical puzzle. Trends in Microbiology. 13, 285–293

10. De Graaf, M., Van Beek, J., Vennema, H., Podkolzin, A. T., Hewitt, J., Bucardo, F., Templeton, K., Mans, J., Nordgren, J., Reuter, G., Lynch, M., Rasmussen, L. D., Iritani, N., Chan, M. C., Martella, V., Ambert-Balay, K., Vinjé, J., White, P. A., and Koopmans, M. P. (2015) Emergence of a novel GII.17 norovirus – End of the GII.4 era? Eurosurveillance. 10.2807/1560-7917.ES2015.20.26.21178

11. Chan, M. C. W., Lee, N., Hung, T.-N., Kwok, K., Cheung, K., Tin, E. K. Y., Lai, R. W. M., Nelson, E. A. S., Leung, T. F., and Chan, P. K. S. (2015) Rapid emergence and predominance of a broadly recognizing and fast-evolving norovirus GII.17 variant in late 2014. Nat Commun. 6, 10061

12. Pletneva, M. A., Sosnovtsev, S. V., and Green, K. Y. (2001) The Genome of Hawaii Virus and its Relationship with other Members of the caliciviridae. Virus Genes. 23, 5–16

13. Du, J., Gu, Q., Liu, Y., Li, Q., Guo, T., and Liu, Y. (2021) The endemic GII.4 norovirus-like-particle induced-antibody lacks of cross-reactivity against the epidemic GII.17 strain. Journal of Medical Virology. 93, 3974–3979

14. Dai, Y.-C., Xia, M., Huang, Q., Tan, M., Qin, L., Zhuang, Y.-L., Long, Y., Li, J.-D., Jiang, X., and Zhang, X.-F. (2017) Characterization of Antigenic Relatedness between GII.4 and GII.17 Noroviruses by Use of Serum Samples from Norovirus-Infected Patients. J Clin Microbiol. 55, 3366–3373

15. Vongpunsawad, S., Venkataram Prasad, B. V., and Estes, M. K. (2013) Norwalk Virus Minor Capsid Protein VP2 Associates within the VP1 Shell Domain. J Virol. 87, 4818–4825

16. Prasad, B. V. V., Hardy, M. E., Dokland, T., Bella, J., Rossmann, M. G., and Estes, M. K. (1999) X-ray Crystallographic Structure of the Norwalk Virus Capsid. Science. 286, 287–290

17. Czakó, R., Atmar, R. L., Opekun, A. R., Gilger, M. A., Graham, D. Y., and Estes, M. K. (2015) Experimental Human Infection with Norwalk Virus Elicits a Surrogate Neutralizing Antibody Response with Cross-Genogroup Activity. Clin. Vaccine Immunol. 22, 221–228

18. Lindesmith, L., Moe, C., LePendu, J., Frelinger, J. A., Treanor, J., and Baric, R. S. (2005) Cellular and Humoral Immunity following Snow Mountain Virus Challenge. J Virol. 79, 2900–2909

19. Lindesmith, L. C., McDaniel, J. R., Changela, A., Verardi, R., Kerr, S. A., Costantini, V., Brewer-Jensen, P. D., Mallory, M. L., Voss, W. N., Boutz, D. R., Blazeck, J. J., Ippolito, G. C., Vinje, J., Kwong, P. D., Georgiou, G., and Baric, R. S. (2019) Sera Antibody Repertoire Analyses Reveal Mechanisms of Broad and Pandemic Strain Neutralizing Responses after Human Norovirus Vaccination. Immunity. 50, 1530–1541.e8

20. Alvarado, G., Ettayebi, K., Atmar, R. L., Bombardi, R. G., Kose, N., Estes, M. K., and Crowe, J. E. (2018) Human Monoclonal Antibodies That Neutralize Pandemic GII.4 Noroviruses. Gastroenterology. 155, 1898–1907

21. Sapparapu, G., Czakó, R., Alvarado, G., Shanker, S., Prasad, B. V. V., Atmar, R. L., Estes, M. K., and Crowe, J. E. (2016) Frequent Use of the IgA Isotype in Human B Cells Encoding Potent Norovirus-Specific Monoclonal Antibodies That Block HBGA Binding. PLoS Pathog. 12, e1005719

22. Tanaka, T., Kitamoto, N., Jiang, X., and Estes, M. K. (2006) High Efficiency Cross-Reactive Monoclonal Antibody Production by Oral Immunization with Recombinant Norwalk Virus-Like Particles. Microbiology and Immunology. 50, 883–888

23. Gray, J. J., Cunliffe, C., Ball, J., Graham, D. Y., Desselberger, U., and Estes, M. K. (1994) Detection of immunoglobulin M (IgM), IgA, and IgG Norwalk virus-specific antibodies by indirect enzyme-linked immunosorbent assay with baculovirus-expressed Norwalk virus capsid antigen in adult volunteers challenged with Norwalk virus. J Clin Microbiol. 32, 3059–3063

24. Van Loben Sels, J. M., and Green, K. Y. (2019) The Antigenic Topology of Norovirus as Defined by B and T Cell Epitope Mapping: Implications for Universal Vaccines and Therapeutics. Viruses. 11, 432

25. Lindesmith, L. C., Beltramello, M., Donaldson, E. F., Corti, D., Swanstrom, J., Debbink, K., Lanzavecchia, A., and Baric, R. S. (2012) Immunogenetic Mechanisms Driving Norovirus GII.4 Antigenic Variation. PLoS Pathog. 8, e1002705

26. Parra, G. I., Azure, J., Fischer, R., Bok, K., Sandoval-Jaime, C., Sosnovtsev, S. V., Sander, P., and Green, K. Y. (2013) Identification of a Broadly Cross-Reactive Epitope in the Inner Shell of the Norovirus Capsid. PLoS ONE. 8, e67592

27. Alvarado, G., Salmen, W., Ettayebi, K., Hu, L., Sankaran, B., Estes, M. K., Venkataram Prasad, B. V., and Crowe, J. E. (2021) Broadly cross-reactive human antibodies that inhibit genogroup I and II noroviruses. Nat Commun. 12, 4320

28. Yi, Y., Wang, X., Wang, S., Xiong, P., Liu, Q., Zhang, C., Yin, F., and Huang, Z. (2021) Identification of a blockade epitope of human norovirus GII.17. Emerging Microbes & Infections. 10, 954–963

29. Strother, C. A., Brewer-Jensen, P. D., Becker-Dreps, S., Zepeda, O., May, S., Gonzalez, F., Reyes, Y., McElvany, B. D., Averill, A. M., Mallory, M. L., Montmayeur, A. M., Costantini, V. P., Vinjé, J., Baric, R. S., Bucardo, F., Lindesmith, L. C., and Diehl, S. A. (2023) Infant antibody and B-cell responses following confirmed pediatric GII.17 norovirus infections functionally distinguish GII.17 genetic clusters. Front. Immunol. 14, 1229724

30. Keyt, B. A., Baliga, R., Sinclair, A. M., Carroll, S. F., and Peterson, M. S. (2020) Structure, Function, and Therapeutic Use of IgM Antibodies. Antibodies. 9, 53

31. Wibroe, P. P., Helvig, S. Y., and Moein Moghimi, S. (2014) The Role of Complement in Antibody Therapy for Infectious Diseases. Microbiol Spectr. 2, 2.2.10

32. Oostindie, S. C., Lazar, G. A., Schuurman, J., and Parren, P. W. H. I. (2022) Avidity in antibody effector functions and biotherapeutic drug design. Nat Rev Drug Discov. 21, 715–735

